# Hyper-*k*-mers: efficient streaming *k*-mers representation

**DOI:** 10.1101/2024.11.06.620789

**Authors:** Igor Martayan, Lucas Robidou, Yoshihiro Shibuya, Antoine Limasset

**Author notes:** co-first authors (Igor Martayan, Lucas Robidou).

## Abstract

*K*-mers have become ubiquitous in modern bioinformatics pipelines. A key factor in their success is the ability to filter out erroneous *k*-mers by removing those with low abundances. However, large numbers of distinct *k*-mers make counting a memory-intensive step. Early tools addressed this issue by storing *k*-mers on disk. More recent solutions mitigate the excessive redundancy of overlapping *k*-mers by partially reassembling them into super-*k*-mers. Nevertheless, consecutive super-*k*-mers still overlap by *k* − 1 bases, leading to some degree of inefficiency.

Here we present *hyper-k-mers* as an alternative, less redundant, representation of super-*k*-mers. Our contributions are three-fold. First, we propose hyper-*k*-mers, a new *k*-mer representation that asymptotically decreases duplication compared to super-*k*-mers. Second, we present a theoretical analysis comparing the space efficiency of super-*k*-mers, syncmers, and hyper-*k*-mers. Our approach offers significant advantages compared to super-*k*-mers, by reducing the asymptotic lower bound from 6 to 4 bits per nucleotide. Third, we present `KFC`, a *k*-mer counting algorithm leveraging hyper-*k*-mers. `KFC` offers significant practical advantages, including an order of magnitude improvement in memory usage compared to state-of-the-art tools. Notably, our experiments show that `KFC` is the only tool whose memory usage scales sub-linearly with *k*-mer size *k*, and is the fastest option when *k* is large.

**Availability:** `KFC` is available at https://github.com/lrobidou/KFC with tests available at https://github.com/imartayan/KFC_experiments.

## 1 Introduction

Representing DNA sequences as sets of *k*-mers, fixed-length sub-strings of length *k*, is a simple yet highly efficient approach in sequence bioinformatics by enabling accurate estimation of sequence similarity in linear time, circumventing the need for computationally intensive alignment steps. Moreover, it naturally handles redundancy, as each unique *k*-mer is stored only once, regardless of its frequency in the DNA sequence. This property is particularly beneficial when dealing with sequencing datasets that are inherently redundant due to high coverage depths (in the order of 1000x in large sequencing datasets). Over the years, the use of *k*-mers has been extended to various applications such as genotyping [15], phylogeny reconstruction [40], and read error correction [19]. However, naive storage of *k*-mers leads to significant space overheads, potentially up to a factor of *k*, since each position in the DNA sequence can generate a *k*-mer. Additionally, sequencing errors introduce erroneous non-biologically-relevant *k*-mers. As observed by Velvet [43], for sufficiently large *k*’s, spurious *k*-mers only repeat a few times, since it is unlikely for two independent errors to produce the same *k*-mer. However, due to high coverage depths, the total number of erroneous *k*-mers can easily outnumber the true genomic *k*-mers by orders of magnitude. Filtering out infrequent *k*-mers, while retaining the highly frequent ones, has proven to be an extremely effective strategy for distinguishing true *k*-mers from errors. This observation led to one of the simplest, yet most fundamental tasks in computational biology, known as *k-mer counting*, which aims to associate *k*-mers to their occurrence counts in a dataset for filtering purposes. Despite its apparent simplicity, *k*-mer counting is quite resource-intensive, as storing large numbers of *k*-mers along with their counts is expensive in terms of both memory and time. Given their critical role in various applications, *k*-mer counting tools have historically focused on speed while limiting memory usage to practical levels. A significant milestone was achieved with Jellyfish [24], which implements a mutex-free hash table for high-throughput parallelism, achieving counting speeds akin to simple file reading. Alternative approaches have utilized more efficient data structures for counting, such as counting quotient filters [28,27], Burst tries [20], and cuckoo hashing [42]. However, these methods still face the challenge of storing entire *k*-mers in main memory. Some tools trade results accuracy for reduced memory usage by using probabilistic data structures like Bloom filters [25,34] or quotient filters [28] to remove low abundance *k*-mers before counting. Another approach to reduce memory is to partition *k*-mers into buckets. Buckets can be safely stored on disk and independently processed thus limiting the number of *k*-mers in main memory [7,31]. Finally, some methods have attempted to accelerate *k*-mer counting by leveraging GPUs [41,11].

A significant advancement in reducing the memory footprint of *k*-mer storage was the introduction of super-*k*-mers [18,7]. Super-*k*-mers encode *N* successive *k*-mers sharing the same minimizer – the smallest sub-string of length *m* within a *k*-mer, for a given ordering [35,32,3] – into *k*+*N*− 1 nucleotides. Minimizers exhibit locality-sensitive hashing properties, functioning as a special case of MinHash with a single fingerprint, and have been widely used in *k*-mer partitioning [4,23]. With the behavior of minimizers now well understood [33], we can accurately estimate the representation cost of super-*k*-mers, leading to significant improvements in efficiency. The adoption of super-*k*-mers has mitigated the memory issues of *k*-mer counting, and super-*k*-mer-based tools such as KMC3 [16] and FASTK [26] are among the best tools of this competitive field.

Even in the absence of errors, efficient *k*-mer representations are still required to minimize memory footprint. However, super-*k*-mers are still far from the optimal space efficiency, due to overlaps at their extremities, leading to unwanted redundancies. Over the years, multiple efficient encodings of *k*-mer sets have been proposed [3,36,37], and allowed lightweight *k*-mer based tools [5,29,12]. The common underlying principle is that redundancy can be decreased by merging overlapping *k*-mers. One of the most well-known instances of this idea is the concept of *unitigs*, which are sequences formed by non-branching paths in de Bruijn graphs [3]. Unitigs are one of the most well-known de Bruijn graph representations, with duplications only occurring near branching paths. Subsequent representation proposed to allow more merging even if they may be unsure biologically to be more succinct. Such representations are essentially static, as they are constructed once from a complete set of *k*-mers. In contrast, super-*k*-mers can be constructed on-the-fly during sequence parsing, making them a more versatile representation. Moreover, this representation is inherently partitioned, since each super-*k*-mer (and thus each *k*-mer) is associated with its minimizer, which naturally facilitates indexing [21,22,30]. A fundamental question is whether we can improve this convenient light-weight and partitioned representation to be as memory-efficient as static ones, thereby obtaining the best of both worlds. Static representations achieve space close to the lower bound of 2 bits per nucleotide (over the four letters DNA alphabet), whereas super-*k*-mer representations are several times larger for commonly-used *k*-mer sizes (*k* ≈ 31). However, for *k* →+∞, super-*k*-mers tend to 6 bits per *k*-mer [33]. Up until now, *k*-mer sizes have been quite small, due to the limited read lengths of next-generation sequencing (NGS) datasets. However, larger *k*-mer sizes are becoming more practical thanks to continuous improvements in the accuracy of long-read sequencing technologies, such as Oxford Nanopore Technologies (ONT), whose reads are approaching 1% error rates [38], and PacBio HiFi sequencing with error rates as small as 0.1% [1]. These advancements enable the use of very large *k*-mer sizes in applications like genome assembly, *k*-mer analysis, and pangenomics. Conversely, very large *k*-mer sizes degrade the performance of traditional efficient encoding methods (unitigs, simplitigs, eulertigs…), as overlap sizes increase linearly with *k*. To address this challenge, we introduce the concept of *hyper-k-mers* as an alternative to super-*k*-mers, aiming to bridge the gap between dynamic and static representations. Compared to super-*k*-mers, hyper-*k*-mers-based representations are 33% lighter while displaying similar properties. We demonstrate the practical interest of our approach by implementing and testing a new *k*-mer counting tool called `KFC` based on hyper-*k*-mers. `KFC` is both the fastest and the more memory-efficient tool for large *k*’s (*k* ≥ 200), being the only one being more efficient as *k* increases. `KFC` supports KFF, a new standardized format for representing *k*-mer sets [9].

The rest of the paper is organized as follows. In section 2, we formally define the concepts of *k*-mers, minimizers, super-*k*-mers, hyper-*k*-mers, and the main goal of our work. Section 3 presents the theoretical analysis of hyper-*k*-mers and details the algorithm and ideas behind our tool `KFC`. Section 4 reports experimental results to assess `KFC`’s performance compared to other *k*-mer counting tools. We conclude with discussions on possible future directions and general considerations.

## 2 Definitions and problem statement

In this section, we introduce the concepts at the basis of our exposition about hyper-*k*-mers in section 3. For the most part, we re-use the notation from [14] and [29], which we report here for convenience. We considered strings defined over the alphabet *Σ* = {*A, C, G, T*} of size *σ* = 4, and we use the terms *character, base* and *nucleotide* interchangeably. Let *S* ∈ *Σ*^*^ be a (genomic) string. We refer to *S*[*i*..*j*) as the substring of length *j* − *i*, starting at position *i* (included) and ending at position *j* (excluded).

### Definition 1

**(*k*-mer)**. *k-mers are substrings S*[*i*..*i* + *k*) *of fixed length k*.

### Definition 2

**(Order)**. *An order* 𝒪_*m*_ *on m-mers is an injective function* 𝒪_*m*_ : *σ*^*m*^ → R, *such that x* ≤𝒪_*m*_ *y if and only if* 𝒪_*m*_(*x*) ≤ 𝒪_*m*_(*y*)

In practice, (pseudo-)random hash functions *h* : *Σ*^*m*^ → *U* are used to define an order [30], where *U* is a sufficiently large universe of possible values, like *U* = 2^64^ if using 64-bit hash functions.

### Definition 3

**((Random) minimizer)**. *A minimizer of a k-mer K is its minimal substring of length m < k for a given (random) order* 𝒪_*m*_.

Minimizers have been widely studied in the literature [35,32,14] and multiple selection methods, called *minimizer schemes*, have been proposed. The efficiency of a minimizer scheme is usually measured by its *density*.

### Definition 4

**(Density of a minimizer scheme)**. *The density of a minimizer scheme is the expected proportion of selected minimizers over the number of m-mers in a random string*.

In particular, the density of random minimizers is equal to *d* = 2*/*(*w* + 1) [35], where *w* = *k* − *m* + 1 is the window size (the number of *m*-mers in a *k*-mer). Thus, two neighboring *k*-mers overlapping by *k*− 1 bases often share the same minimizer.

### Definition 5

**(Super-*k*-mer)**. *A maximal sequence of consecutive k-mers having the same minimizer* [17].

### Definition 6

**(Overlap of consecutive super-*k*-mers)**. *Given two consecutive super-k-mers u and v, we define the overlap of u and v, denoted* ov(*u, v*), *as the k*− 1 *bases shared by the rightmost k-mer of u and the leftmost k-mer of v*.

A straightforward method to reduce *k*-mer memory consumption is to store super-*k*-mers instead of *k*-mers. Let ℐ be the set of *k*-mers of string *S*. Each character (base) in *S* can be encoded using 2 bits since *σ* = 4. Thus, naively storing set ℐ requires 2*k* bits per nucleotide in *S* (each character in *S* is covered by *k k*-mers, except the extremities), whereas a super-*k*-mer representation only requires 6 bits per nucleotide [33]. While undoubtedly an improvement, super-*k*-mers space usage is still far away from the optimal achievable space of 2 bits per nucleotide of a completely (theoretical) linear representation of ℐ. The remaining overhead comes from the *k* − 1 bases overlaps between neighboring super-*k*-mers.

Our main goal is to find a representation of ℐ whose space is *<* 6 bits per nucleotide.

## 3 Methods

We start this section by introducing theoretical analysis of super-*k*-mers and their space requirements. Next, we do the same for closed syncmers [10]. Syncmers have been shown to “pave” sequences better than other sampling techniques [10,39], with a tendency to cover more bases than, e.g. minimizer-based sampling. Despite not being suitable for representing *k*-mer sets, we think our theoretical results on closed syncmers are useful to put hyper-*k*-mer into perspective.

### 3.1 Super-*k*-mers

#### Lemma 1

**(Average number of super-*k*-mers)**. *Given a random string S, the expected number of super-k-mers is* (|*S*| − *m* + 1) × *d*.

*Proof*. By definition of the density *d*, the expected number of minimizers is (|*S* |−*m* + 1) ×*d*. Since each minimizer corresponds to one super-*k*-mer, the expected number of super-*k*-mers is the same.

#### Lemma 2

**(Average length of super-*k*-mers)**. *Given a random string S, the average length of a super-k-mer is* 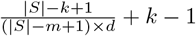.*In particular, it approaches* 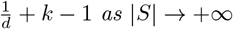.

*Proof*. By Lemma 1, there are (|*S*| − *m* + 1) × *d* super-*k*-mers on average. Since there are |*S*| − *k* + 1 *k*-mers in *S*, each super-*k*-mer contains 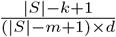 *k*-mers on average, i.e.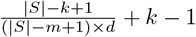.bases.

#### Theorem 1

**(Super-*k*-mer space usage)**. *Given a random string S, the expected total number of bases in the super-k-mers of S is asymptotically equivalent to* |*S*| · (1 + (*k* − 1) · *d*) *as* |*S*| → +∞.

*Proof*. Let *S*_*sk*_ denote the set of super-*k*-mers of *S*, |*S*_*sk*_| the number of super-*k*-mers in *S* and *S*_*sk*_[*i*] denotes the *i*-th super-*k*-mer of *S*. The total number of characters in *S*_*sk*_ is:

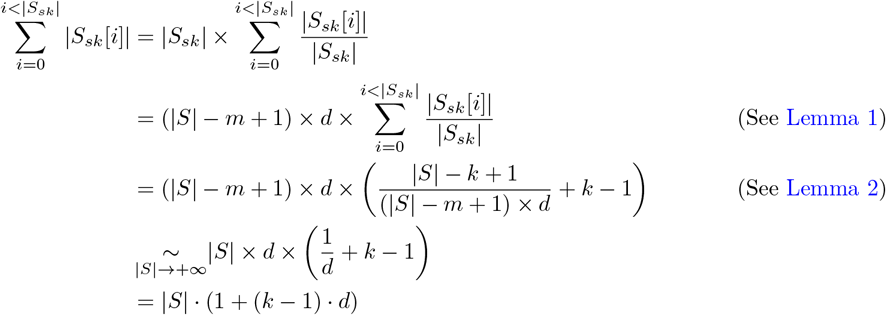

In particular, for random minimizers which have a density *d* = 2*/*(*w* + 1) =(2*/*(*k* − *m* +) 2), the average number of bases in a set of super-*k*-mers is asymptotically equivalent to 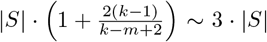 when *k* ≫*m*. Thus, a super-*k*-mer scheme requires 3 bases per nucleotide in |*S* |, i.e. 6 bits per nucleotide in |*S*| for an alphabet of size 4.

### 3.2 Closed-syncmers

#### Definition 7

**(Closed syncmer)**. *A k-mer with a minimizer located at its start or its end*.

#### Theorem 2

**(Closed syncmer space usage)**. *Given a random string S, the expected total number of bases in the closed syncmers of S is asymptotically equivalent to* |*S*| · *k* · *d as* |*S*| → +∞.

*Proof*. Each *k*-mer consists of *w* = *k* − *m* + 1 *m*-mers. Let us consider the last *w* + 1 *m*-mers after spawning a new minimizer. Since a new minimizer has just been found, the smallest *m*-mer, among the last *w* + 1, is either the first (if the minimizer changed because the previous one went out of scope) or the last (if the new minimizer is smaller than the previous one). If the smallest *m*-mer is the first one, then the previous *k*-mer is a closed syncmer. Otherwise, the smallest *m*-mer is at the end of the last *k*-mer, which is thus a closed syncmer. Thus, except the first minimizer, each minimizer creates a closed syncmer, so there are (|*S*| − *m* + 1) × *d* closed syncmers, on average, in *S*. Since each closed syncmer has length *k*, the expected total number of bases is (|*S*| − *m* + 1) × *d* × *k*, which is asymptotically equivalent to |*S*| · *k* · *d* as |*S*| → +∞.

In particular, for random minimizers, the number of bases in a set of closed syncmers is asymptotically equivalent to 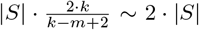,for *k* ≫*m*. As such, closed syncmers require 2 bases per nucleotide in |*S*|, i.e. 4 bits for an alphabet of size 4.

### 3.3 Hyper-*k*-mers

#### Definition 8

**(Hyper-*k*-mer)**. *Given three consecutive super-k-mers u, v and w, let m*_*v*_ *be the minimizer of v. We define the hyper-k-mer associated to m*_*v*_ *as* (ov(*u, v*), ov(*v, w*)), *and refer to* ov(*u, v*) *as its left part and* ov(*v, w*) *as its right part. By convention, if v has no predecessor in the sequence, we define its left part as the first k* − 1 *bases and if it has no successor, we define its right part as the last k* − 1 *bases*.

We can already make a few observations based on this definition: every hyper-*k*-mer contains exactly 2(*k* − 1) bases and given two consecutive hyper-*k*-mers *x* and *y*, the right part of *x* and the left part of *y* are identical. This means that we can effectively save *k*− 1 bases for each pair of consecutive hyper-*k*-mers. Figure 1 gives an example of hyper-*k*-mers and shows that their overlapping parts can be shared.

**Fig. 1:**
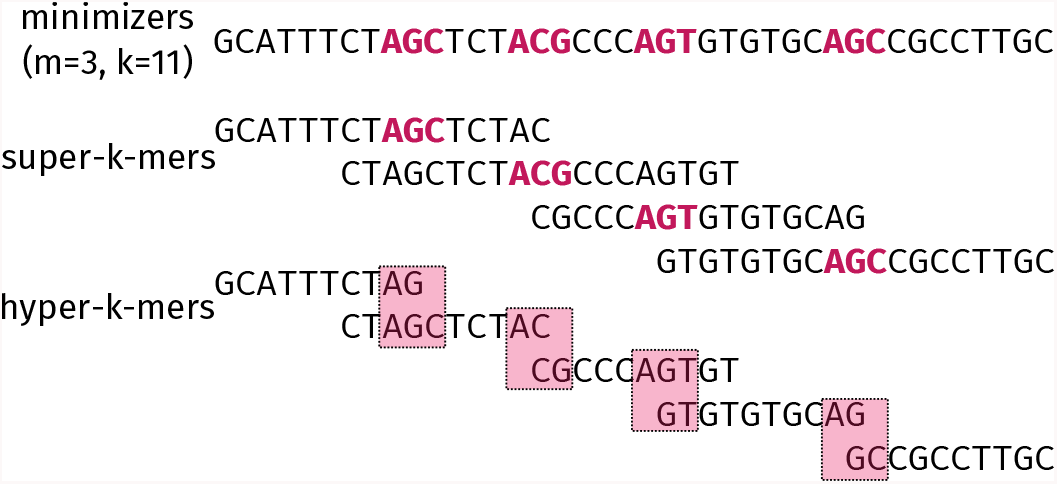
Example of minimizers, super-*k*-mers and hyper-*k*-mers for *m* = 3 and *k* = 11 using lexicographic order. In this example, the original sequence contains 40 bases and super-*k*-mers use 70 bases while hyper-*k*-mers use 50 bases.

#### Theorem 3

**(Hyper-*k*-mer space usage)**

*Given a random string S, the expected total number of bases in the hyper-k-mers of S is asymptotically equivalent to* |*S*| · (*k* − 1) · *d as* |*S*| → +∞. *In addition, storing the links between the left and right parts results in a* 2 log_2_(*d*|*S*|) *bits overhead for each hyper-k-mer*.

*Proof*. Each minimizer of *S* is associated with one hyper-*k*-mer, so there are (|*S*| − *m* + 1) × *d* hyper-*k*-mers on average. What’s more, each pair of consecutive hyper-*k*-mers shares *k* − 1 bases which can be written only once. Thus, except for the first hyper-*k*-mer (which uses 2(*k* − 1) bases), each hyper-*k*-mer uses *k* − 1 bases. Therefore, the expected total number of bases is ((|*S*| − *m* + 1) × *d* + 1) × (*k* − 1), which is asymptotically equivalent to |*S*| · (*k* − 1) · *d* as |*S*| → +∞. Finally, we need to maintain a “link” between the two parts of each hyper-*k*-mer by storing their indices. Since the expected number of hyper-*k*-mers is equivalent to *d*|*S*|, each index takes log_2_(*d*|*S*|) bits of space, leading to an overhead of 2 log_2_(*d*|*S*|) bits for each hyper-*k*-mer.

In particular, for random minimizers, the number of bases in a set if hyper-*k*-mers is asymptotically equivalent to 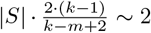 bases per nucleotide in |*S*|, for *k* ≫ *m*. Moreover, assuming *m* = Ω(log_2_ (*d*|*S*|), the space overhead of the links becomes negligible as the space usage as *k* ≫ *m*. Figure 2 gives an overview of the evolution of *k* increases, and compares it to the evolution of super-*k*-mer’s space usage.

**Fig. 2:**
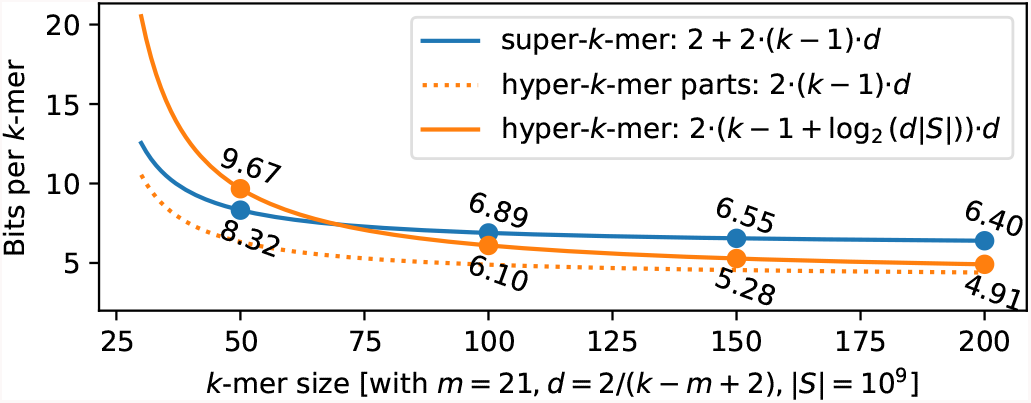
Space usage of super-*k*-mers and hyper-*k*-mers for *k* ∈ [30, 200], with random minimizers of size 21.

These results, summarized in Table 1, show that representing a set of *k*-mers using hyper-*k*-mers is more space-efficient than super-*k*-mers and slightly more space-efficient than closed syncmers, regardless of the chosen minimizer scheme. In particular, hyper-*k*-mers require ^2^/_3_ of the space used by super-*k*-mers for random minimizers, and this ratio gets even lower for minimizer schemes with a lower density.

**Table 1:**
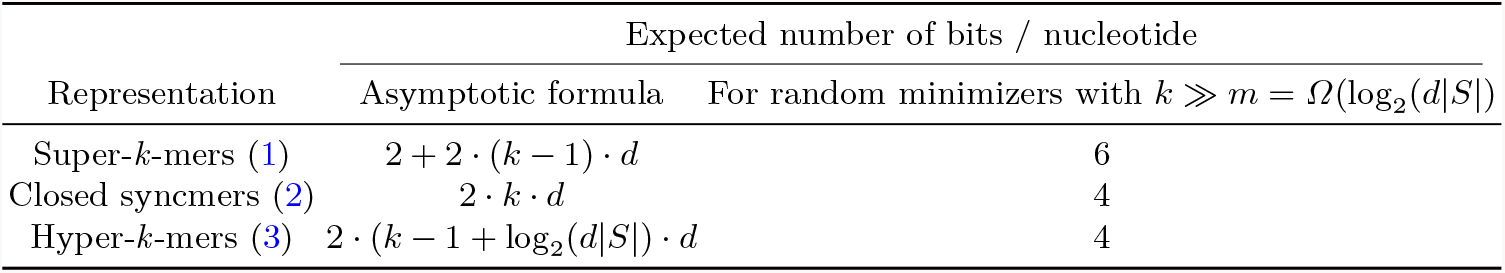
Comparison of the space usage of super-*k*-mers, closed syncmers and hyper-*k*-mers.

### 3.4 Counting *k*-mers using hyper-*k*-mers

This section proposes *k*-mer counting as an application of hyper-*k*-mers to a real-life use case. Hyper-*k*-mers are used as a proxy for storing *k*-mer counts. A simplified overview of the resulting index is presented in Figure 3.

**Fig. 3:**
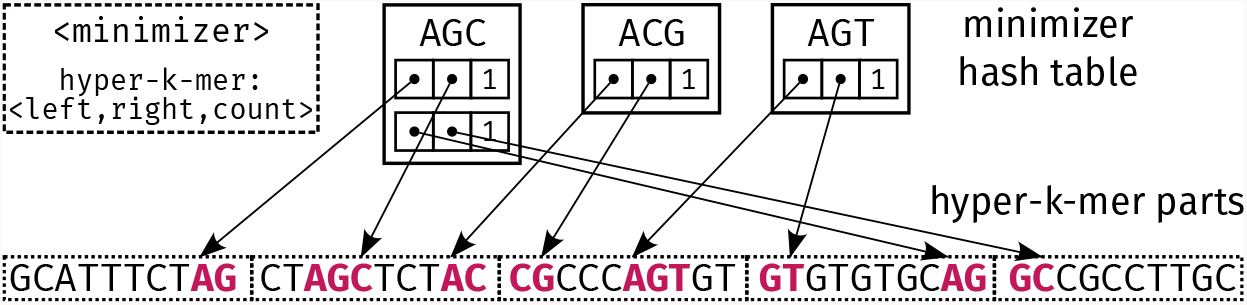
High-level view of the `KFC` data structure, using the example introduced in Figure 1. Two data structures are maintained. **Bottom**: a vector of hyper-*k*-mers parts stores the left and right parts of the hyper-*k*-mers computed from the original set of sequences. The bold parts correspond to the (possibly truncated) minimizers of the sequence. **Top**: a hash table associates each minimizer to a list of entries. Each entry represents a hyper-*k*-mer and its count. Each hyper-*k*-mer is represented using pointers (represented as dots) to its left and right parts in the vector of hyper-*k*-mers parts. To retrieve the parts, each pointer consists of the part’s position in the vector, a start and end position in that part, and an orientation flag to handle reverse complements.

Our *k*-mer counter implements a two-pass algorithm. In the first stage, we iterate over super-*k*-mers in the input sequences. We identify which super-*k*-mers are solid, i.e. are present more than twice in the input. If a super-*k*-mer is solid, the hyper-*k*-mer, along with its minimizer, is computed using its left and right super-*k*-mer neighbors (see Definition 8). If a computed hyper-*k*-mer was already present in the index, we increase its count in the hash table, otherwise, we insert a new entry in the table and insert the corresponding hyper-*k*-mers in the vector (encoded using two bits per base). However, doing so prevents us from counting the first and last super-*k*-mers of a read, as they are likely truncated and, by extension, unique and below our threshold. It also prevents us from counting the non-solid super-*k*-mers sharing their minimizer with a solid super-*k*-mer (indicating a possible sequencing error in the non-solid super-*k*-mer, whose *k*-mers shared with the solid super-*k*-mers should *not* be discarded).

As such, the second stage consists of correcting these issues by re-reading the input. Non-solid super-*k*-mers sharing their minimizer with solid super-*k*-mers which are substrings of hyper-*k*-mers already in the index, are added to the internal hash table. This also allows for the addition of the first and last super-*k*-mers of each read. After these two stages, the only *k*-mers that will be wrongly discarded are the ones that **A/** are present more than two times in the input but **B/** belong to a non-solid super-*k*-mer that does not share its minimizer with any of the solid super-*k*-mers. This super-*k*-mer-level threshold is slightly more aggressive than a *k*-mer-level threshold to filter unique *k*-mers. The effects of this heuristic are discussed in section 4.2.

As our algorithm works in a streaming fashion, at any time, only the index has to be maintained in memory whereas reads are only loaded on-demand. This allows our implementation to have a light memory footprint (see subsection 4.2), and to be parallelized (see Figure 10 in Supplementary Materials).

`KFC`’s algorithm is given in Supplementary Materials (`KFC`’s algorithm). For clarity, we omit some details, such as how `KFC` handles subsequences in hyper-*k*-mers parts.

## 4 Experiments

In this section, we benchmark our proposed *k*-mer counter, `KFC`, based on hyper-*k*-mers and compare it against other counters: Jellyfish [24], KMC [6,7,16], FASTK [26], Kaarme [8] and Gerbil [11]. Our implementation is written in Rust and is publicly available at https://github.com/lrobidou/KFC with accompanying test scripts at https://github.com/imartayan/KFC_experiments.

## 4.1 Experimental setup

We report the datasets used in this study and their characteristics in the Supplementary Materials (Table 2). The experiments were performed on a common laptop computer (Dell Inc. Precision 7780 with 13th Gen Intel^®^ Core i9-13950HX Œ 32 and 64 GB RAM).

**Table 2:** Datasets used for testing KFC. Min. Avg. and Max. refer to the minimum, average and maximum length of the reads, respectively.

| Name | Organism | Type | Coverage | # reads | Total length | Min. | Avg. | Max. |
| --- | --- | --- | --- | --- | --- | --- | --- | --- |
| SRR11434954 | E.Coli | HiFi | 5000x | 1,789,131 | 23,122,913,014 | 46 | 12,924.1 | 26,294 |
| SRR28370642 | E.Coli | ONT (Duplex) | 50x | 114,703 | 236,908,842 | 349 | 2,065.4 | 126,029 |
| SRR28370651 | E.Coli | ONT (Duplex) | 50x | 109,061 | 226,502,819 | 330 | 2,076.8 | 125,012 |
| SRR28370668 | E.Coli | ONT (Simplex) | 2000x | 6,819,683 | 9,101,103,830 | 1 | 1,334.5 | 396,011 |

## 4.2 Results

### Long reads whole genome sequencing datasets

In our initial set of experiments, we evaluate all tools using coverage around 100X of the *E. coli* genome with three distinct long-read methods: ONT duplex, ONT simplex, and HiFi. We tested a wide range of *k*-mer sizes, up to 1021, and present the results in Figure 4.

**Fig. 4:**
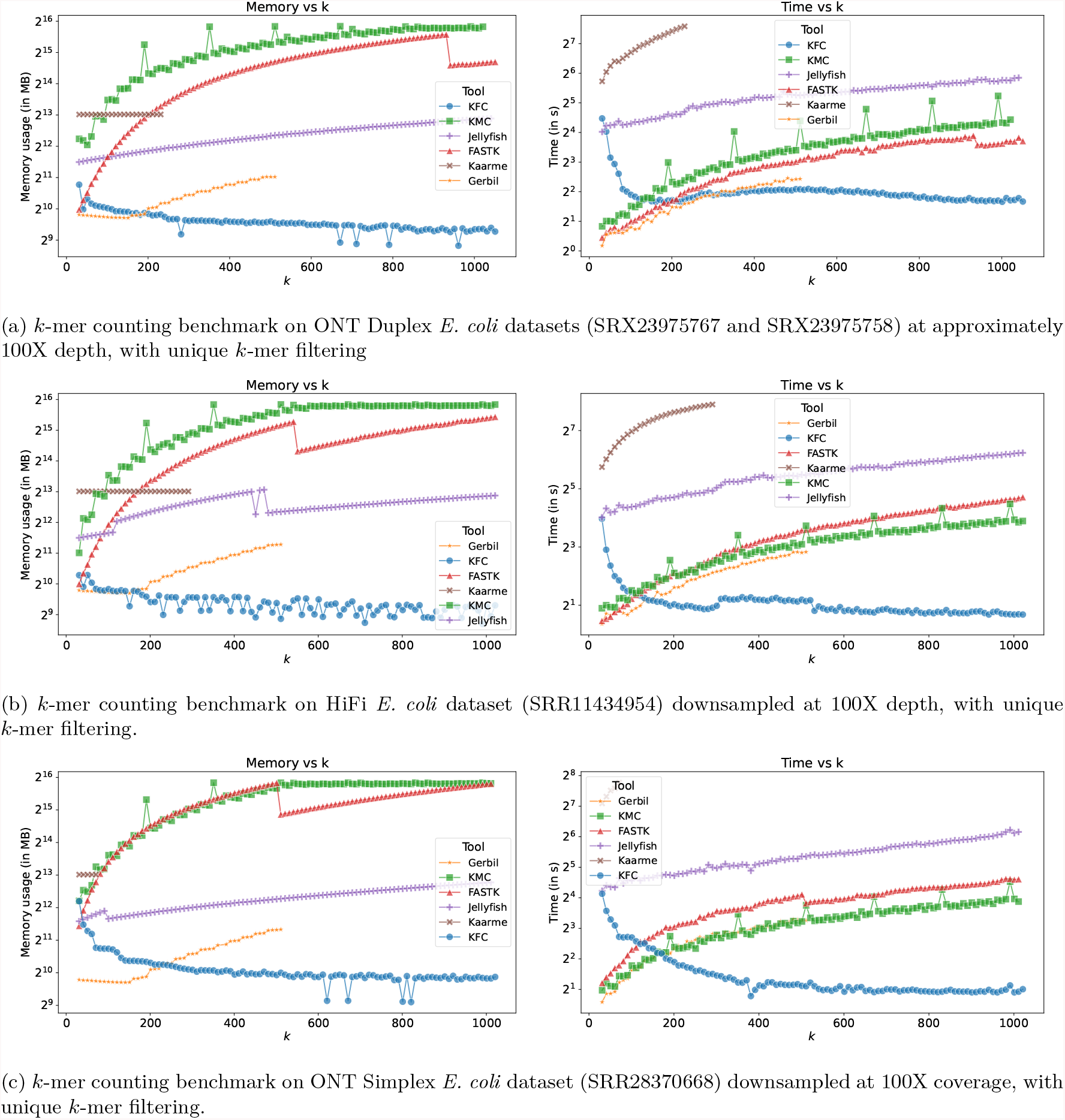
Comparison of *k*-mer benchmarks on different *E*.*coli* datasets. Each subfigure shows the memory usage and timing plots for different sequencing technologies.

Despite the varying error distributions among the methods, the tools exhibit similar overall behaviors. Kaarme maintains an almost constant memory usage but is significantly slower than the other tools. Due to timeouts, it is not tested across all *k*-mer sizes. Jellyfish shows a steadily increasing memory usage with *k*, outperforming FastK and KMC at larger *k*-mer sizes, although it is nearly always the second slowest in terms of runtime. KMC3 and FastK demonstrate very similar behaviors, with memory usage quickly approaching the available RAM. FastK uses less memory than KMC in low-error experiments, but has comparable memory consumption on the noisier simplex dataset. In terms of runtime, both tools behave similarly, with execution time increasing rapidly with *k*. Gerbil outperforms both FastK and KMC3 by using less memory and achieving faster runtimes but cannot handle *k*-mers larger than 512. `KFC` is unique in that it is the only tool to display both decreasing runtime and memory usage with increasing *k*. Consequently, for *k*-mer sizes above 200, `KFC` exhibits the lowest runtime and memory usage among all the tested tools. Experiments for different coverages are reported in the Supplementary material, without significant change in the tools’ respective behaviors. We ran the same benchmark on two larger-scale metagenomic HiFi experiments, in order to assess the tools’ behavior on larger, and more complex datasets. As a medium-scale experiment, we chose a HiFi Zymo Biomics community sequencing that comprises at least one order magnitude more distinct *k*-mers, as it contains 21 strains of 17 species. We sampled 5 Gigabases of accession SRR13128014 and report the results in Figure 5. As a large-scale experiment, we chose the human gut metagenome HiFi sequencings used as metagenome assembly benchmark [13,2] accessions SRR15275210, SRR15275211, SRR15275212, and SRR15275213. Pooled, assemblers estimate the genome content of these datasets between 700 and 900 megabases [13]. We sampled 15 Gigabases of the four datasets pooled and report the results in Figure 6. Tests for the full dataset are shown in Figure 12 of Supplementary materials.

**Fig. 5:**
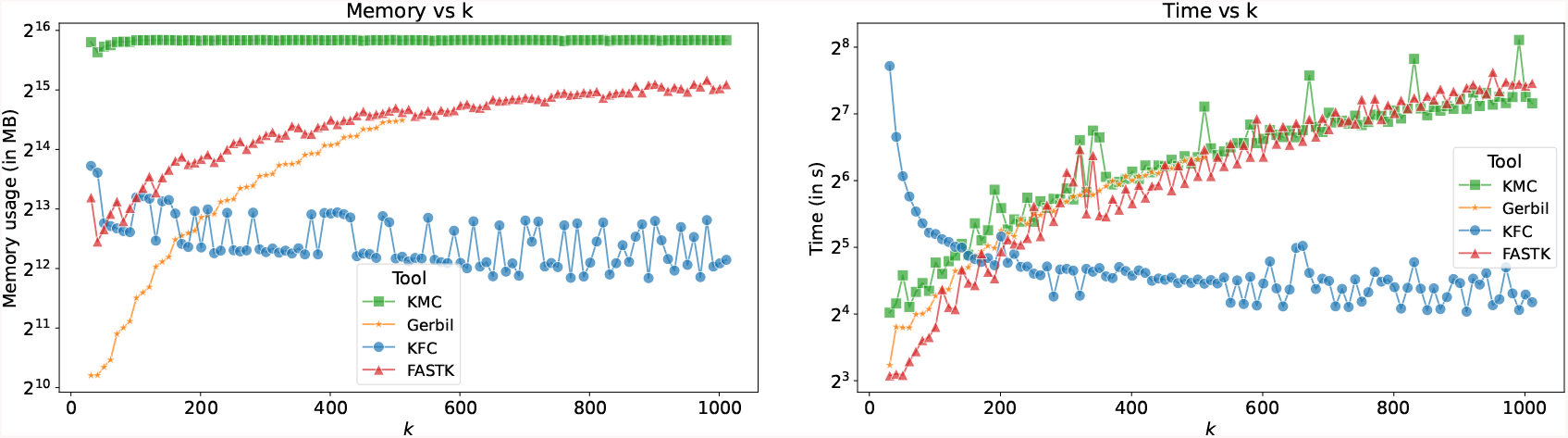
*k*-mer counting benchmark on HiFi Zymo community dataset (SRR13128014) downsampled at 5 gigabases, with unique *k*-mer filtering.

**Fig. 6:**
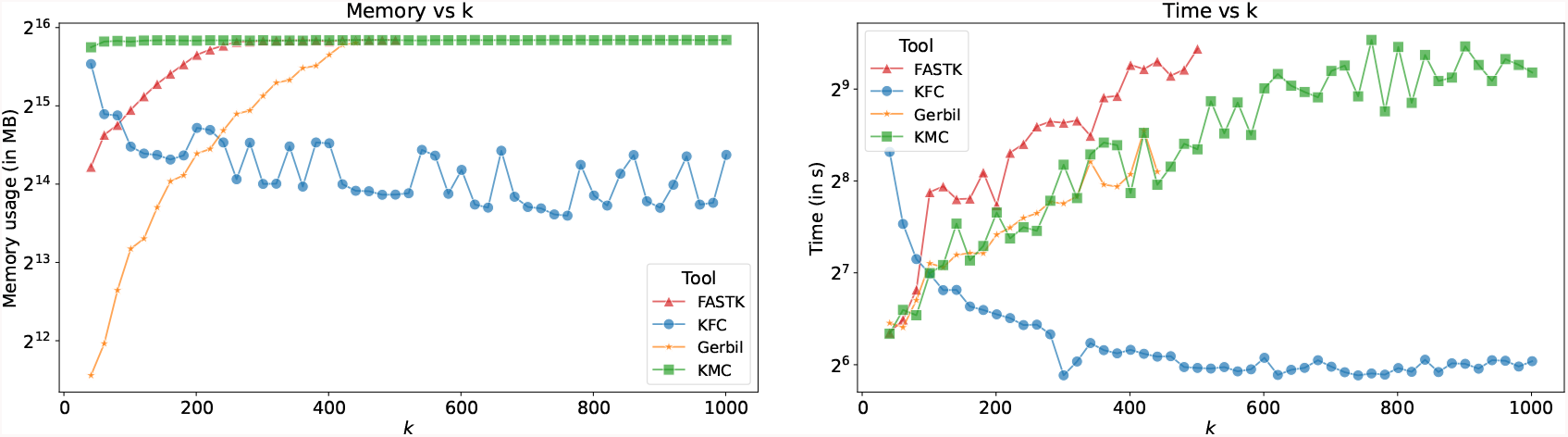
*k*-mer counting benchmark on HiFi human gut datasets (SRR15275210,SRR15275211,SRR15275212,SRR15275213) downsampled at 15 gigabases, with unique *k*-mer filtering.

One might question how much of the observed improvement is due to the loss of certain *k*-mers resulting from the super-*k*-mer filtering heuristic. To confirm the validity of our approach, we conduct the same experiment without unique *k*-mer filtering, i.e., counting all *k*-mers and therefore without super-*k*-mer filtering. The results are shown in Supplementary materials (Figure 13). We observe that Jellyfish’s performance remains unchanged, while FastK and KMC’s performance significantly degrades in both time and memory usage. While `KFC` is indeed slower and more memory-intensive without the super-*k*-mer filtering heuristic, it remains competitive even when compared to its competitors that filter unique *k*-mers.

### Pangenome *k*-mer analysis

In a distinct experiment reported in Figure 7, we run the different tools on one thousand *S. enterica* complete genomes from NCBI to evaluate the cost of counting large *k*-mers for pangenome analysis. We observe that `KFC` remains the most memory-efficient tool, followed by FastK and then KMC. Regarding runtime, `KFC` becomes the fastest tool for very large *k*-mer sizes (above 500), despite its currently limited multithreading efficiency compared to KMC when processing low amounts of sequences. This suggests the need for further software engineering improvements for `KFC`, as it is not yet mature. Notably, when using a lower number of threads, `KFC` becomes the fastest tool. Additionally, FastK’s runtime increases rapidly with k. These findings highlight the advantage of `KFC` in handling genome collections without filtering.

**Fig. 7:**
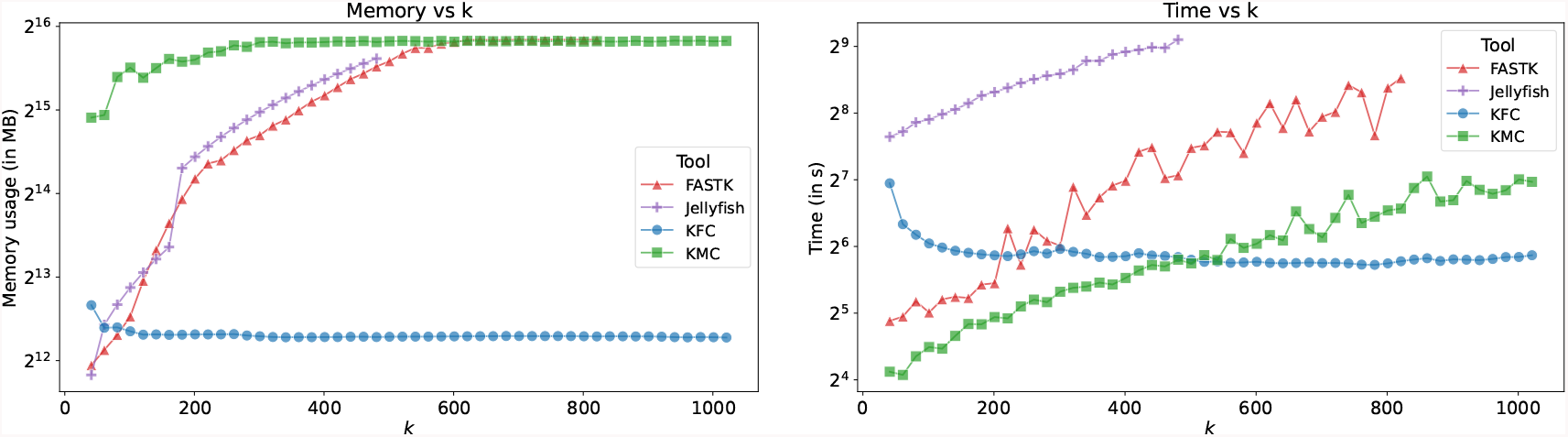
*k*-mer counting benchmark on one thousand *S. enterica* complete genome.

### Internal evaluations

In this section, we present several internal experiments to evaluate key implementation aspects of `KFC`.

We evaluated the super-*k*-mer abundance threshold heuristic by analyzing the number of *k*-mers lost when applying the unique super-*k*-mer removal heuristics (Figure 8). We observe that the number of “lost” *k*-mers decreases exponentially as the *k*-mer abundance threshold increases. This loss is more significant at low coverage levels, particularly for larger *k*-mer sizes. Similar to classical *k*-mer abundance filtering, higher error rates significantly impact the mean coverage of genomic *k*-mers, resulting in greater *k*-mer loss at high error rates compared to low error rates. Although the proportion of lost *k*-mers may seem substantial, we argue that most of these are low-abundance ones and thus likely erroneous. To substantiate this claim, we count the number of *k*-mers from the reference genome that are absent in both KMC and `KFC` outputs, considering different *k*-mer sizes and abundance thresholds (Figure 9). At low *k*-mer abundance thresholds, `KFC` filters more *k*-mers than KMC; however, as the *k*-mer abundance threshold increases, both methods exhibit similar behaviors, with the number of filtered *k*-mers remaining relatively low. We repeated the experiment using a dataset with a HiFi error rate (0.1%) and observed no differences between the outputs of KMC and `KFC`. This confirms that the super-*k*-mer abundance threshold, while slightly more aggressive than the *k*-mer-level threshold, remains conservative enough to maintain accuracy while delivering substantial performance improvements.

**Fig. 8:**
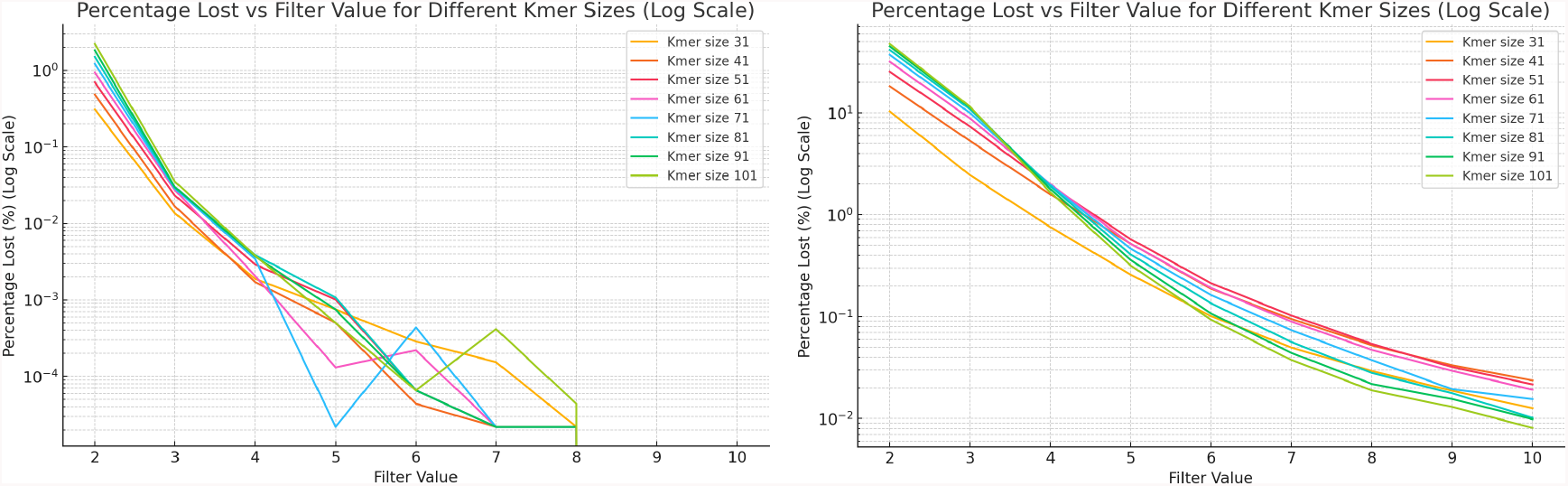
Percentage of *k*-mers lost by removing unique super-*k*-mers, based on the applied abundance threshold and *k*-mer size. The plot on the left represents a simulated HiFi dataset with 100x coverage and an error rate of 0.1%, while the plot on the right shows results from a simulated ONT dataset with the same coverage but an error rate of 1%.

**Fig. 9:**
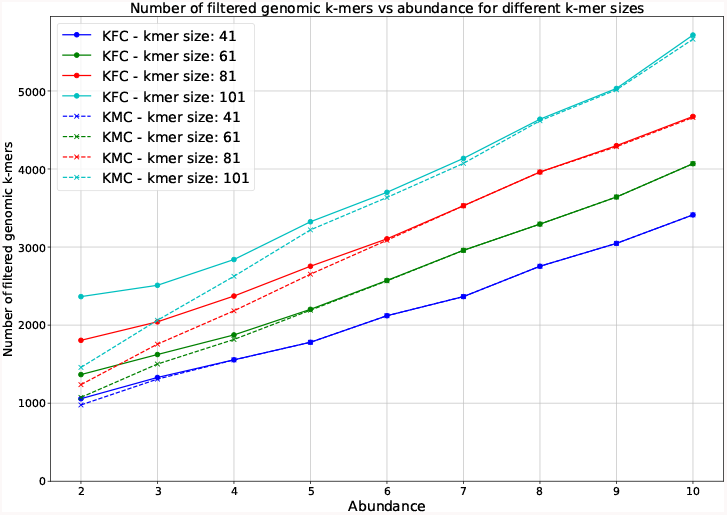
Amount of filtered genomic *k*-mers by `KFC` and KMC according to the abundance threshold and the *k*-mer size on a simulated 100X ONT *E. coli* dataset with 1% error rate.

**Fig. 10:**
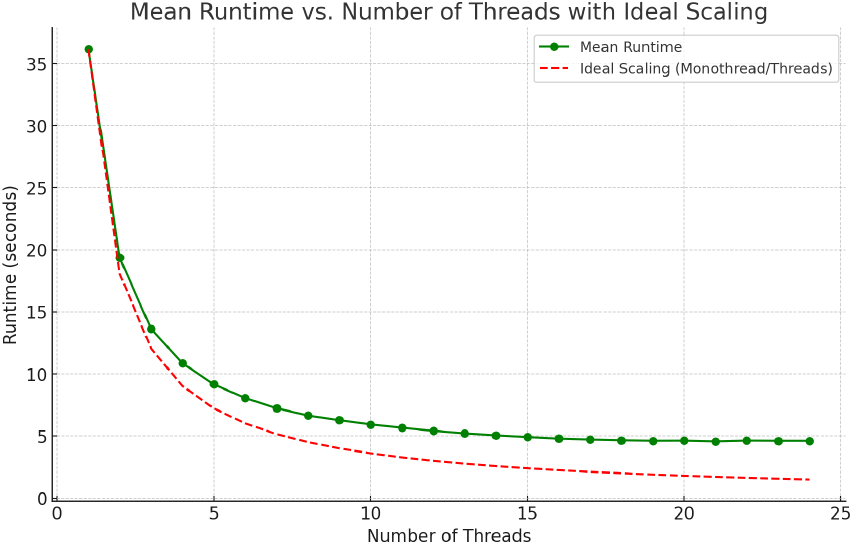
Multiple core usage efficiency on the 100X HiFi *E. coli* dataset.

We also report the efficiency of `KFC`’s multithreading in the Supplementary Materials, subsection A.2.

## 5 Conclusions and future work

In this work, we studied the problem of memory-efficient *k*-mer set representations and proposed a novel solution, *hyper-k-mers*, which aims to reduce space redundancy caused by *k*-mer overlaps. Theoretical analysis and practical experiments demonstrate that hyper-*k*-mers are more succinct than their direct competitor, *super-k-mers*, achieving space usage closer to sampling techniques such as syncmers, while still providing a direct *k*-mer representation. Hyper-*k*-mers represent sequences as collections of minimizers and the sequences between them. This approach reduces *k*-mer overlaps and, consequently, theoretically allows hyper-*k*-mers to achieve a space usage of 4 bits per (DNA) base. We implemented a novel *k*-mer counter, `KFC`, based on hyper-*k*-mers, which shows excellent performance. For large *k* values (e.g. *k* ≥ 200), `KFC` outperforms all competitors in both speed and memory usage, and uniquely becomes more efficient for increasing *k*-mer sizes. Since longer *k*-mers are expected to become more relevant in the coming years—thanks to steadily improving sequencing technologies leading to longer and less error-prone reads—we believe that hyper-*k*-mers and `KFC` are future-proof solutions.

Several improvements are possible. First, although our theoretical analysis shows that hyper-*k*-mers are a direct improvement over super-*k*-mers, they are still more space-consuming than the theoretical lower bound of 2 bits per nucleotide. Finding an optimal representation that achieves this lower bound remains an open question. Second, our current implementation of hyper-*k*-mers does not scale well to short *k* values. Smaller *k*’s are harder to optimize due to the large number of collisions between minimizers and contexts. While very small *k*’s (*k <* 16) are manageable with a simple hash table storing *k*-mers individually, *k*-mer lengths in the range 17 to 63 could still benefit from dynamic linear set representations like hyper-*k*-mers. Developing such a solution remains an open question. Finally, as mentioned in subsection 3.4, the current version of `KFC` is based on a two-pass algorithm that discards *k*-mers belonging to non-solid super-*k*-mers if no minimizer is shared with any solid super-*k*-mer. While this heuristic is extremely efficient, handling this corner case is another direction for future improvements.

## Acknowledgments

This work is partially funded by the French National Research Agency AGATE ANR-21-CE45-0012 and full-RNA ANR-22-CE45-0007. With financial support from ITMO Cancer of Aviesan within the framework of the 2021-2030 Cancer Control Strategy, on funds administered by Inserm. The authors would like to thank Giulio Ermanno Pibiri for his input.

## A. Supplementary materials

### A.1 Datasets characteristics

In this section, we report the datasets we used in our experiments and some of their characteristics (Table 2).

### A.2 Multi-threading efficiency of KFC

In this section, we assess the multi-threading efficiency of `KFC` (Figure 10). Our results demonstrate that `KFC` effectively utilizes multi-core architectures, achieving performance gains with up to several dozen cores before experiencing diminishing returns.

### A.3 Effect of coverage

In this section, we present the performance benchmark similar to Figure 4, but with various coverage. The results of this benchmark are shown in Figure 11.

**Fig. 11:**
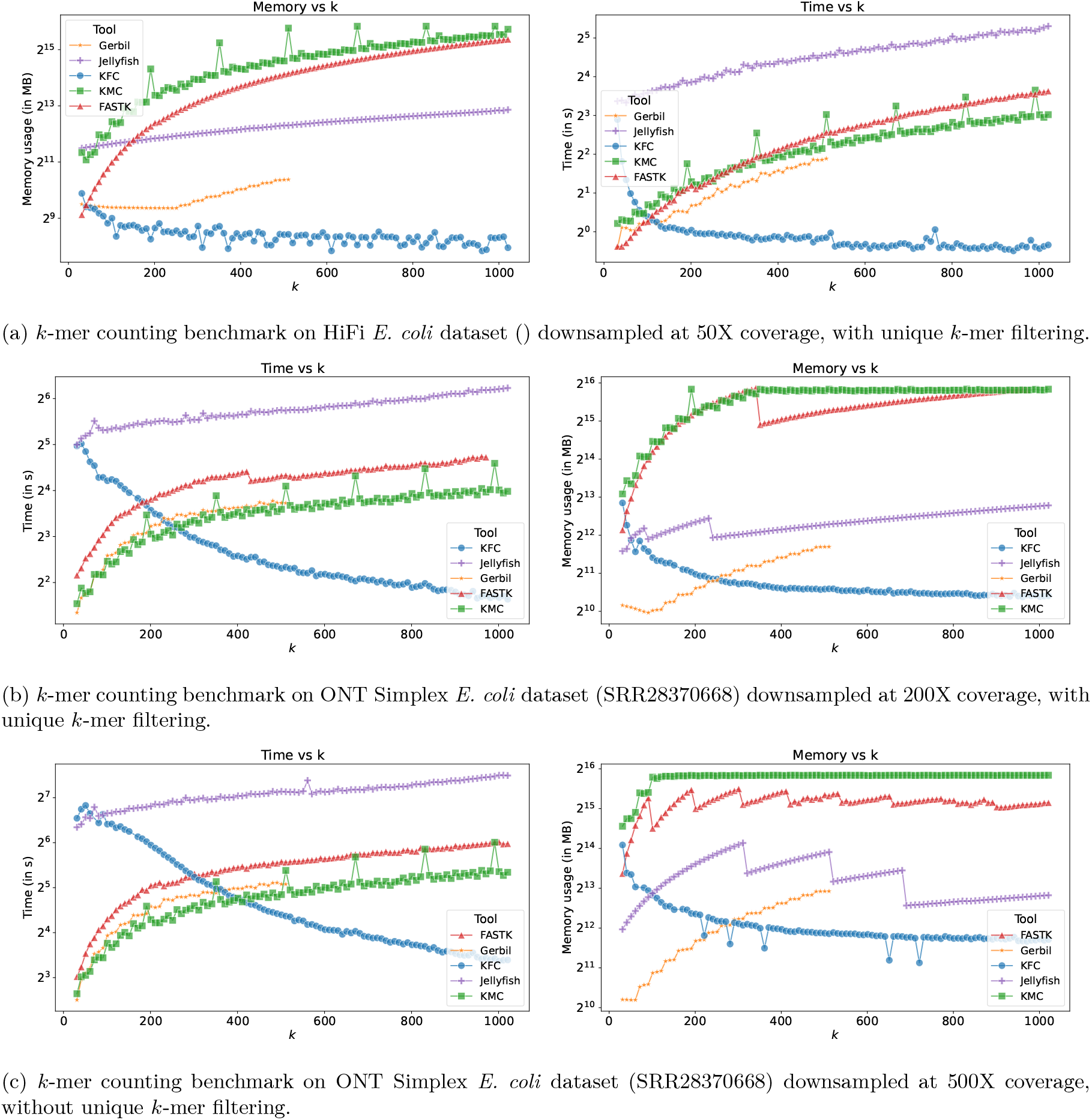
*k*-mer counting benchmark on ONT Simplex *E. coli* dataset (SRR28370668) for various coverage.

**Fig. 12:**
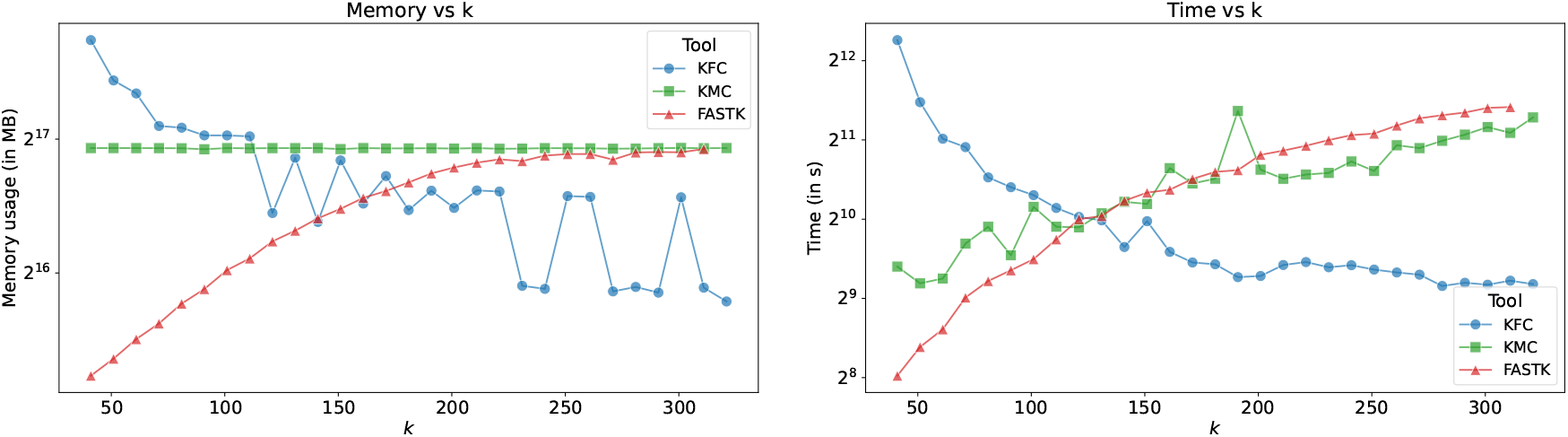
*k*-mer counting benchmark on complete HiFi human gut datasets (SRR15275210,SRR15275211,SRR15275212,SRR15275213), filtering *k*-mers appearing once and twice (threshold equal to 3).

**Fig. 13:**
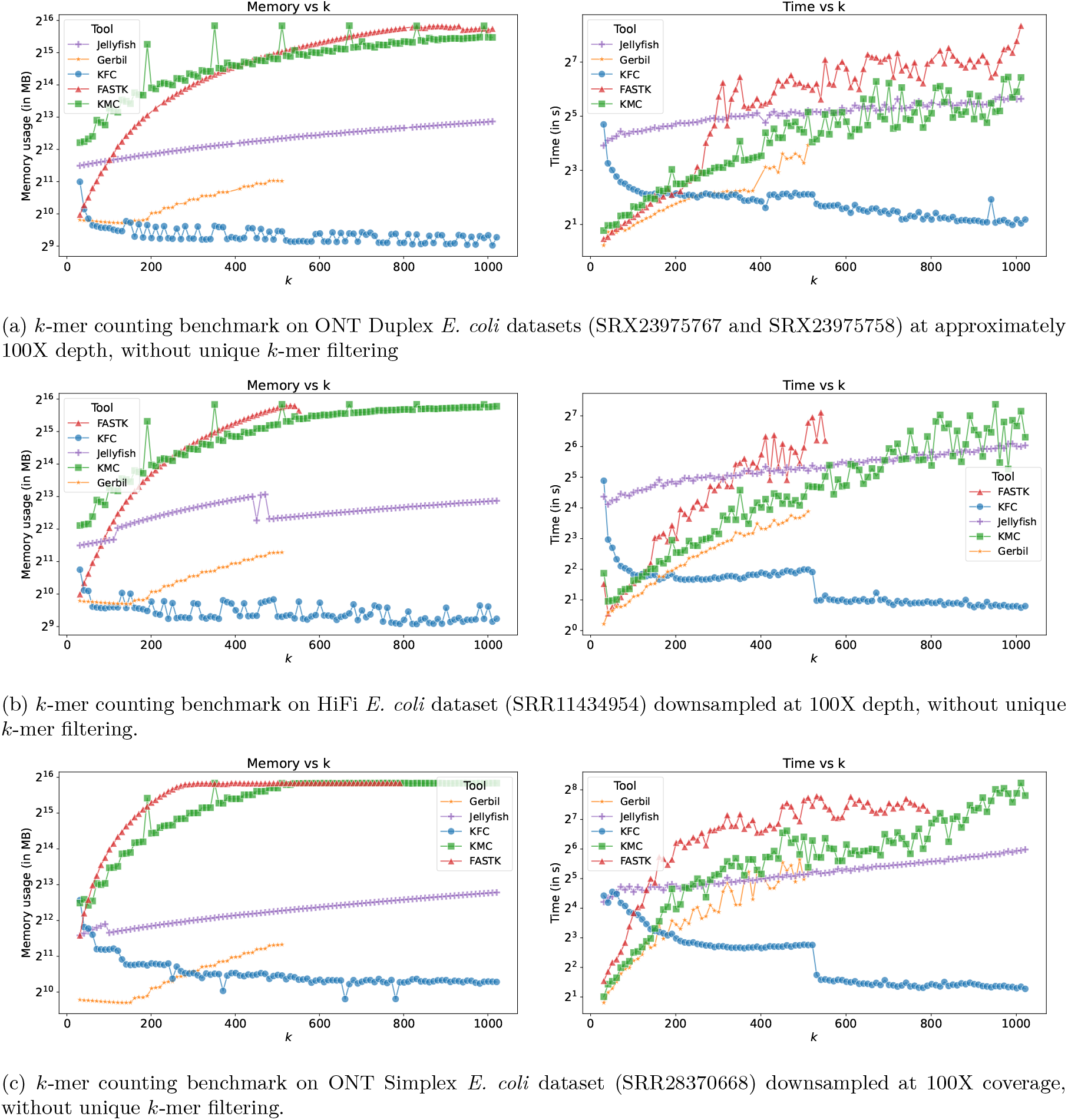
Comparison of Kmer benchmarks on different *E*.*coli* datasets without any filtering. Each subfigure shows the memory usage and timing plots for different sequencing technologies.

### A.4 Effect of not filtering unique *k*-mers

In this section, we present the performance of `KFC` when unique *k*-mers are not filtered out. Figure 13 shows the results discussed in section 4.2. They are similar to Figure 4, but without filtering unique *k*-mers.

### A.5 KFC’s algorithm

In this section, we present an overview of the two stages of `KFC` (in Algorithm 1).

#### Algorithm 1

Counting

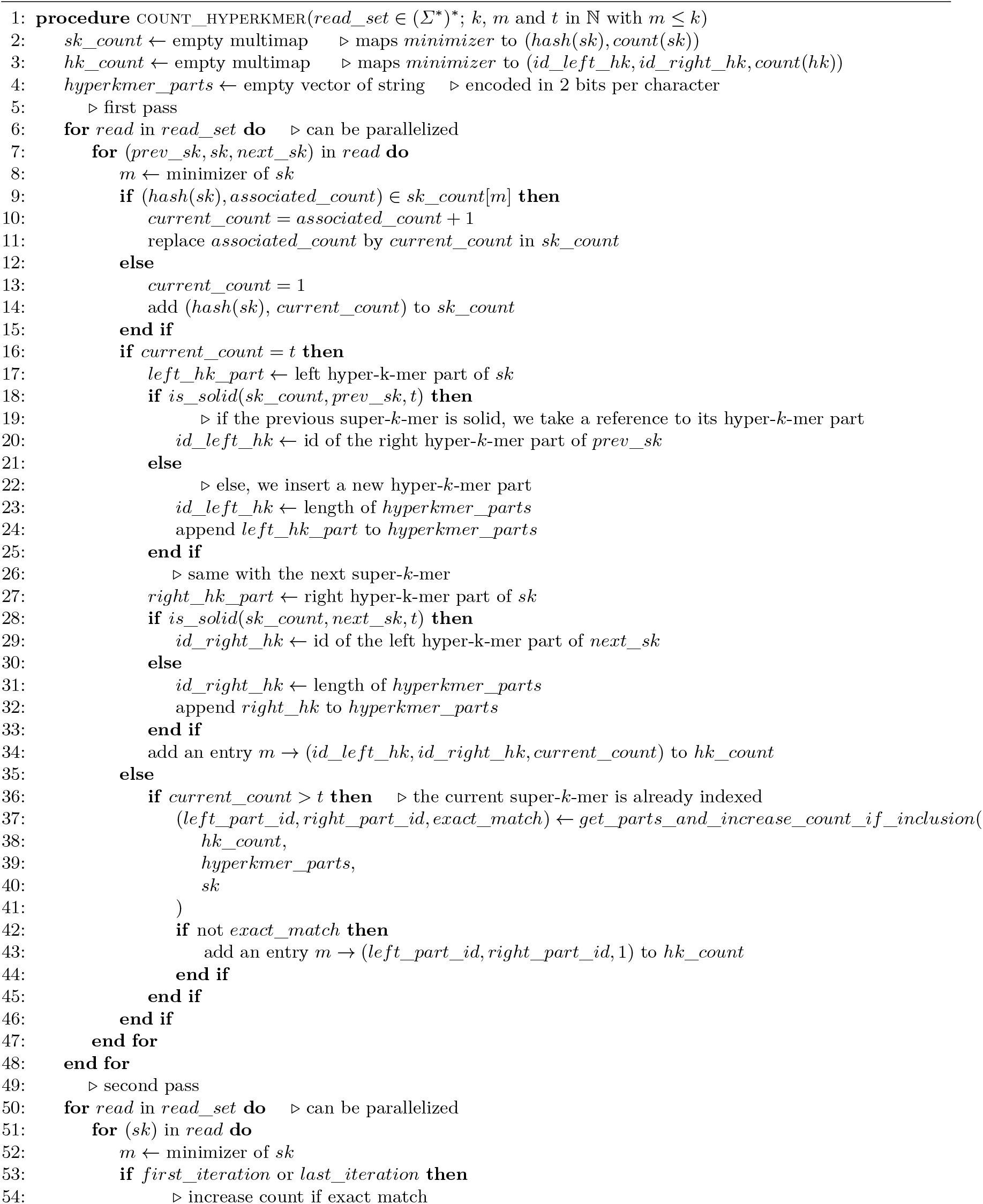
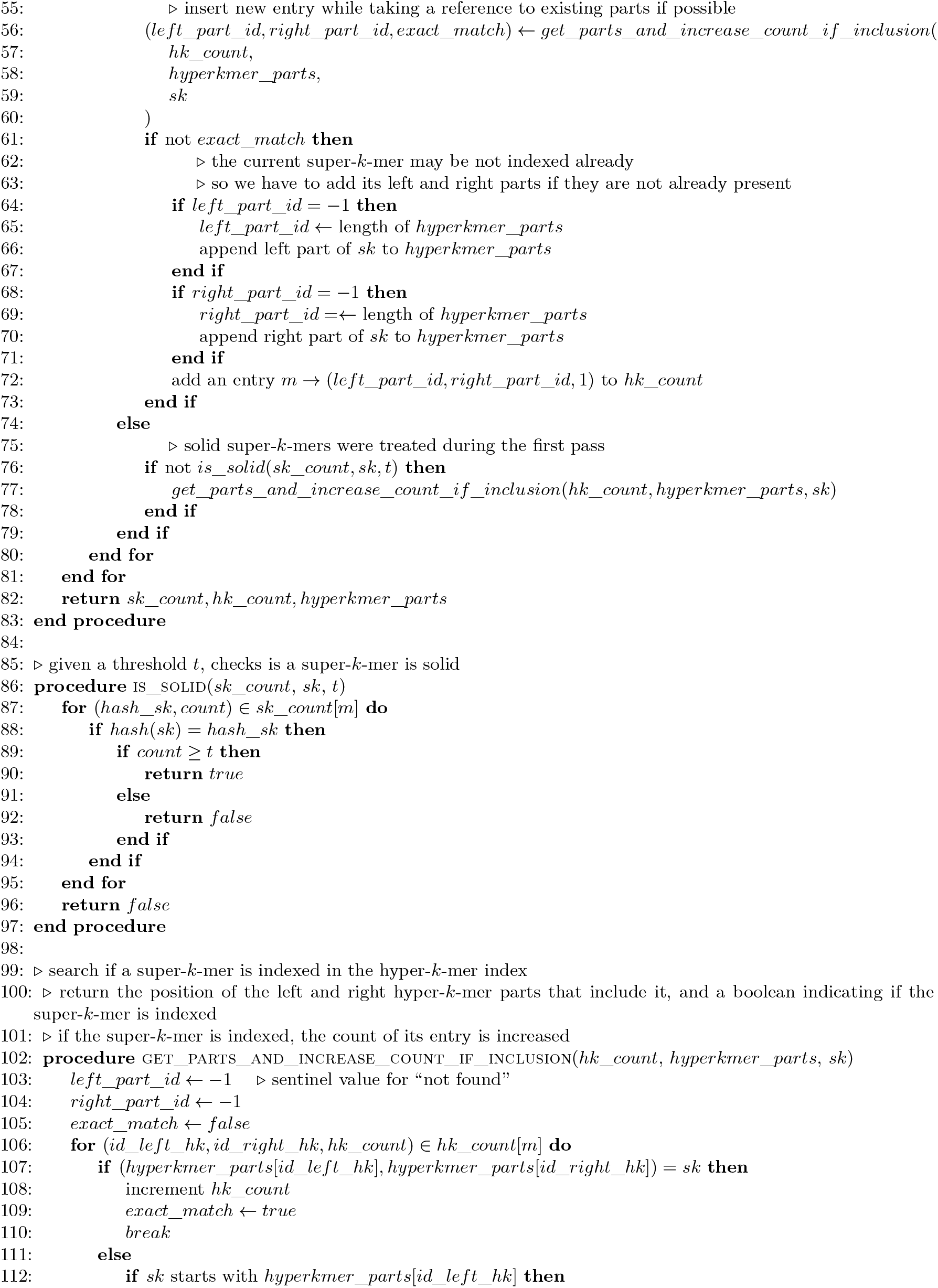
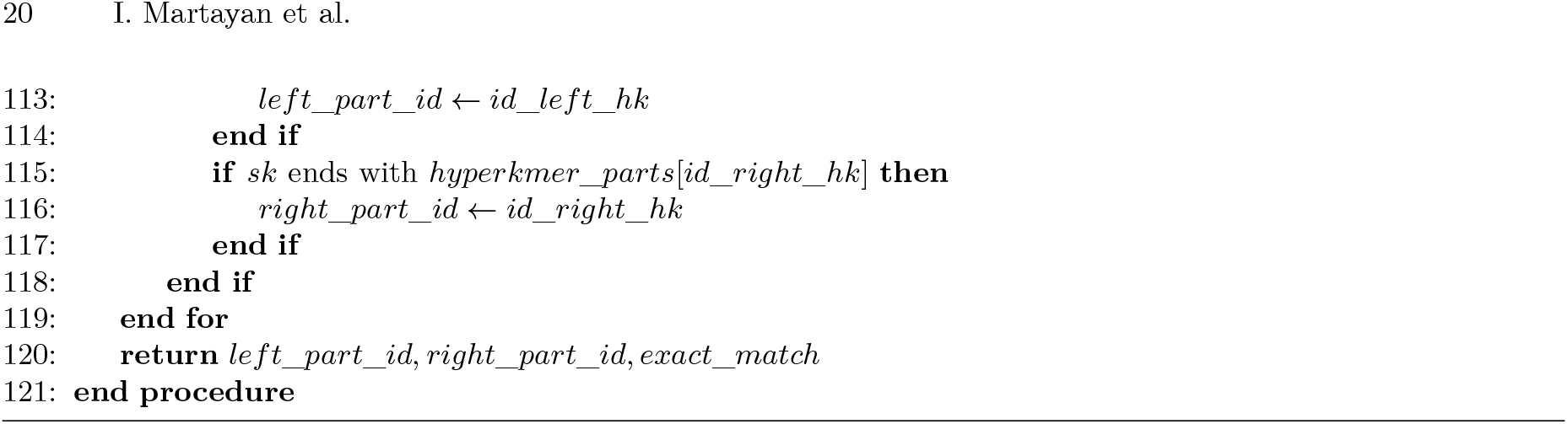

